# Prediction of individual melodic contour processing in sensory association cortices from resting state functional connectivity

**DOI:** 10.1101/2025.03.07.642076

**Authors:** C Ahrends, M Lumaca, ML Kringelbach, D Vidaurre, P Vuust

## Abstract

Recent studies suggest that it is possible to predict an individual brain’s spatial activation pattern in response to a paradigm from their functional connectivity at rest (rsFC). However, it is unclear whether this prediction works across the brain. We here aim to understand whether individual task activation can be best predicted in local regions that are highly specialised to the task at hand or whether there are domain-independent regions in the brain that carry most information about the individual. To answer this question, we used fMRI data from participants (nonmusicians, N=52) at rest and during an auditory oddball paradigm while watching a silent movie. We then predicted individual differences in brain responses to melodic deviants from their rsFC both across the whole brain (global parcellation in 22 regions) and within the auditory cortices (local parcellation in 22 regions). Predictability was consistently higher in specific brain areas. These areas are centred around sensory association cortices: In the local (auditory cortex) parcellation, the best predicted area is the right superior temporal gyrus (STG), an auditory association area. The right STG is a critical region in melodic contour processing, a capacity that is central to the paradigm at hand. Interestingly, the best predicted network in the global parcellation is the bilateral visual association cortex. Sensory association cortices may carry more information specific to an individual regardless of the particular paradigm. Our results indicate that individual differences can be predicted in paradigm-relevant areas or general areas with higher inter-individual variability.

## Introduction

The exact configuration of resting state functional connectivity (rsFC) is unique to an individual, akin to a fingerprint (Biswal et al., 2010; Finn et al., 2015; Horien et al., 2019; Marek et al., 2019). Furthermore, unique patterns that are found in rsFC are also present when an individual is engaged in a task (Finn et al., 2015; Kraus et al., 2021). Individual functional brain networks at rest and during diverse tasks have, in fact, been found to largely overlap, forming a stable functional architecture (Kraus et al., 2021; Krienen et al., 2014; Smith et al., 2009).

Given this stable, unique functional architecture, there have been recent efforts to *predict* what an individual’s brain response related to a paradigm will look like based on their rsFC (Cohen et al., 2020; Cole et al., 2016; Langs et al., 2015; Niu et al., 2021; Osher et al., 2019; Parker Jones et al., 2016; Tavor et al., 2016; Tobyne et al., 2018). For instance, using an fMRI dataset of 98 participants and 47 contrasts derived from 7 different paradigms, Tavor et al. (2016) showed that it is possible to predict individual patterns of task activation with great spatial detail and participant specificity. However, a recent evaluation of different methods attempting this prediction has found that most methods fail to perform better than robust baseline models when predicting across the whole cortex (Lacosse et al., 2021). They also found that, among 47 different task contrasts tested, only one could reliably be predicted. Interestingly, Lacosse et al. (2021) also showed that there are differences between brain areas in terms of how well this activation can be predicted. It is unclear why these differences in predictability between brain areas exist.

One reason may be that certain regions are more relevant for the paradigm at hand. Variation in activation within the relevant area may then simply reflect individual differences in the process studied. Another reason may be that some brain networks show more variability across individuals than others, and that this variability is systematic. The functional and structural organization of these networks would be more sensitive to the influence of idiosyncratic factors (genetics, environment, experience, etc.) and their properties may carry more unique information than others about an individual’s activation patterns and behaviour (Lumaca, Baggio, et al., 2021; Lumaca et al., 2019). In biometrics, fingerprints are used because they are thought to be unique to an individual (Pankanti et al., 2002), while other body parts may not be likely to identify a person unequivocally. This may be similar in the brain, where inter-individual variability may be stronger in some regions than in others.

In the present study, we aim to determine in which parts of the brain rsFC can predict individual differences in melodic processing. To study melodic processing, we employed a paradigm that elicits responses to melodic intervals and melodic contour. Melodic intervals and contour are general features of a musical melody, i.e., a sequence of tones. Melodic intervals refer to the size of a pitch change, i.e., the step or jump size between two subsequent tones. Melodic contour refers to the direction (up or down) in which it progresses without regard to the exact size of pitch change. The ability to distinguish melodic intervals and contour is a fundamental human ability (Dowling & Fujitani, 1971; Peretz & Babaï, 1992) that plays a key role in both language and music perception (Patel 2010). Automatic encoding of melodies is already present in humans in the late foetal stage (Granier-Deferre et al., 2011). Melodic processing varies between individuals based on a number of factors, such as musical expertise (Bailes, 2010; Fujioka et al., 2004; Halpern et al., 1998), specific training (Lo et al., 2015), and age (Jeong & Ryu, 2016). In the brain, melodic processing is associated with activity in the superior temporal gyrus (STG) (Schindler et al., 2013; Stewart et al., 2006) and there is evidence that this activity is predominantly right-lateralised (Johnsrude et al., 2000; Lee et al., 2011; Peretz, 1990) (though, see Schindler et al. 2013 and Stewart et al., 2008 for work challenging a rightward lateralisation). Since melodic processing is spatially relatively confined to the temporal lobe, this makes it a useful paradigm for studying whether paradigm-relevant brain areas, i.e. the STG, or paradigm-unrelated but highly individual brain areas can be better predicted by rsFC.

To study these differences in predictability between brain areas, we used fMRI recordings where participants first rest and then passively listen to a melodic oddball paradigm. Using this dataset, we have previously shown that melodic deviants are mainly associated with activity in subregions of the auditory cortices, namely the STG and Heschl’s gyri (Lumaca, Dietz, et al., 2021). We here created a model to predict each individual’s brain response to the auditory deviants from their rsFC during the previous scanning session. We hypothesised that inter-individual differences in functional activation in our paradigm can be better predicted by rsFC in these subregions of the auditory cortices. To determine these differences in predictability of individual activation patterns between the paradigm-relevant regions and the whole brain, we employed both a whole-brain parcellation and an auditory cortex sub-parcellation.

## Methods

### Study design and data acquisition

Fifty-two healthy adult volunteers (33 females, mean age 24.5 years, range 20–34, all right-handed and non-musicians) participated in the study. MRI data were acquired on a 3T MRI scanner (Siemens Skyra). Participants first completed a functional scan at rest (“resting state data”, 9 mins duration), followed by a structural scan (10 mins duration) and a second functional scan with a paradigm (“paradigm data”, 25 mins duration). During the paradigm scan, participants were presented with an auditory oddball paradigm while watching a subtitled silent documentary. The structural scan consisted of T1-weighted high-resolution imaging sequence using an MP2RAGE sequence (TR = 5,000 ms, TE = 2.87 ms, voxel size = 0.9 mm^3^). Functional images were acquired using a fast multi-band EPI sequence (TR = 1,000 ms, TE = 29.6 ms, voxel size = 2.5 mm^3^). The auditory oddball paradigm was presented via MR-compatible headphones. We presented participants with a stream of melodic patterns: standards (80% frequency), contour deviants (10%), and interval deviants (10%). Each “standard” pattern consisted of 5 different tones drawn from the equal-tempered version of the Bohlen-Pierce scale (BP scale). In the contour “deviants”, the fourth tone violated the surface structure (‘ups’ and ‘downs’) of the sequence, but not the interval size; vice versa for the interval “deviants”. To create variation, patterns were randomly transposed at three different registers of the BP scale (low, medium, and high) (for more details see Lumaca, Dietz, et al. (2021); Lumaca et al. (2019)).

### Pre-processing

We used similar, but distinct pre-processing pipelines for resting state and paradigm data due to the different analysis aims (functional connectivity analysis of resting state data and Generalized Linear Model (GLM)-analysis of paradigm data).

Resting state data was pre-processed in FMRIB’s Software Library (FSL (Jenkinson et al., 2012)). The pipeline consisted of minimal spatial pre-processing, followed by temporal artefact removal. Namely, skull and neck segments were removed from the images using BET brain extraction. The first three volumes of all functional scans were discarded. Initial motion correction (realignment) was carried out using MCFLIRT. We set a threshold of 0.5 mm mean relative rms displacement for possible exclusion of participants due to head motion. No participants exceeded this threshold. The functional scans were then high-pass filtered at 1/2000 Hz and slice-time corrected for the specific slice acquisition order. We used spatial smoothing with a Gaussian kernel of 6 mm FWHM. We then registered all functional scans to their respective structural scans and to MNI space using FLIRT. For temporal artefact removal, each participant’s scanning session (in native space) was decomposed into independent components (ICs) using MELODIC Independent Component Analysis (ICA) (Beckmann, 2012). We trained FIX (Griffanti et al., 2014; Salimi-Khorshidi et al., 2014) to classify ICs into noise and signal based on 10 participants’ hand-labelled session components, which were labelled following guidelines for manual artefact classification (Griffanti et al., 2017). The identified artefactual components, as well as 24 motion regressors were regressed out of each participant’s functional data using unique variance clean-up. The individual functional scans were then transformed to MNI space to perform ICA on the group level.

Pre-processing of the paradigm data is described in detail in Lumaca, Dietz, et al. (2021); Lumaca et al. (2019). Briefly, using SPM12 (r7487), fMRI data were realigned, co-registered, normalised to MNI space, and spatially smoothed (6 mm FWHM). Time series were high-pass filtered at 1/128 Hz and corrected for serial auto-correlations using an AR(1) process, including white noise.

### Data-driven parcellations

The goal of ICA decomposition on the group level is the creation of a data-driven parcellation. Group-level components are voxels which share temporal variance across the time courses of all participants. Each of these group components can be considered a functional parcel. The size and distribution of these components can vary between a small, local cluster of voxels or a distributed network like for instance the default mode network (DMN). We therefore here refer to the components as parcels rather than regions or networks.

To create this group ICA parcellation, the voxel-level resting state data of all participants in MNI space were temporally concatenated. These group time series were then decomposed into ICs using MELODIC ICA (Beckmann et al., 2009). Using this approach, we estimated 25 components across the resting state scanning sessions of all participants. The ICA dimensionality was manually chosen to match typical coarse data-driven functional parcellations, such as the groupICA25 parcellation from the Human Connectome Project (Smith, Beckmann, et al., 2013). Three of these components were removed as (group-level) noise components following visual inspection, resulting in 22 parcels for the final parcellation. The resulting whole-brain (“global”) parcellation is illustrated in **Figure** 1a (top panel). Please note that we here show a binarised version of the parcellation, in which overlap between parcels has been removed because it is simpler to visualise. In the original parcellation, parcel membership of each voxel across the brain is weighted. We use the original non-binarised weighted parcellation to extract resting state time courses and the binarised, non-overlapping version to summarise the paradigm contrast in each region, as explained below.

**Figure 1.**
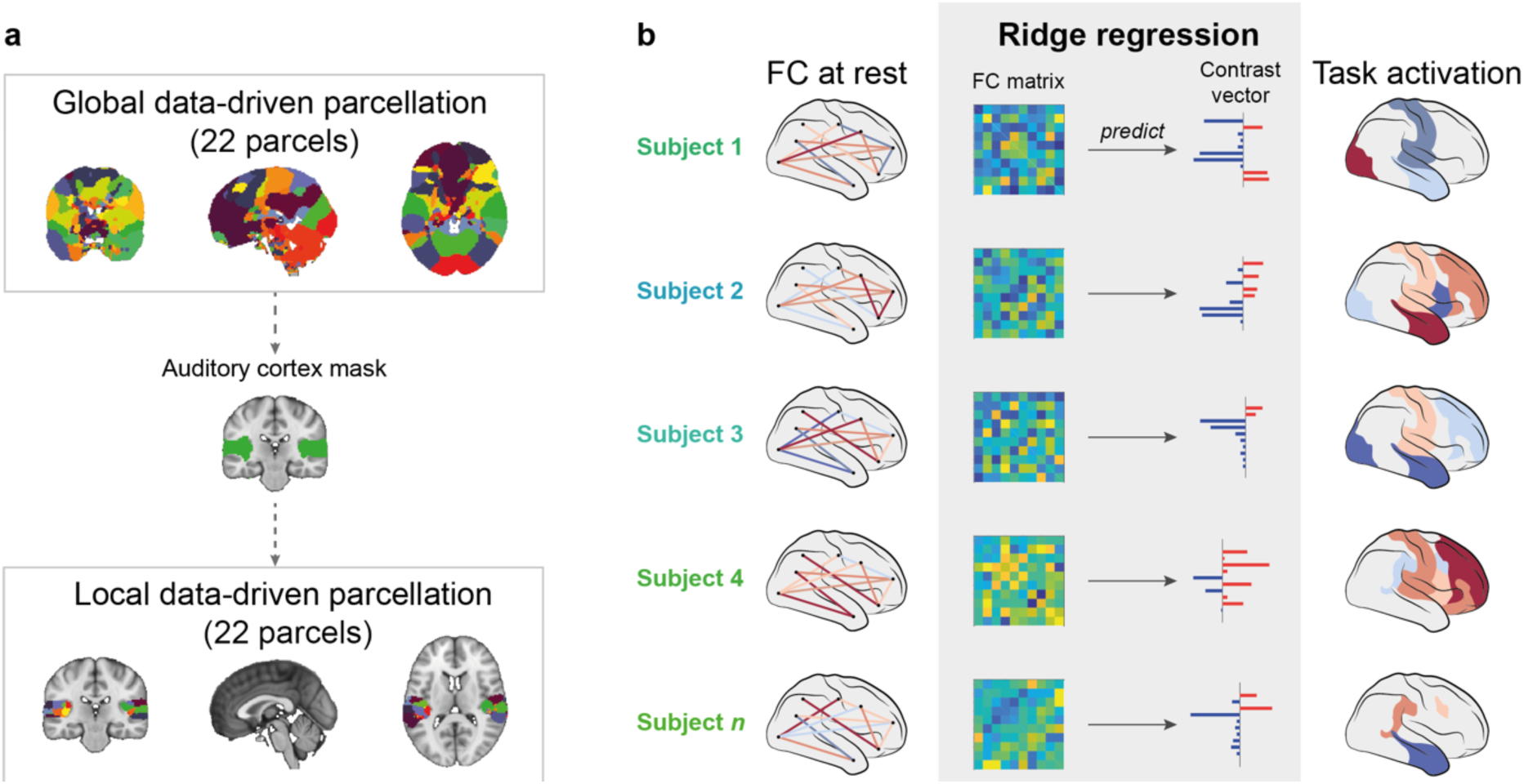
Data-driven parcellations and regression model. **a)** We used two different parcellations to extract time courses of the resting state data and parcellate the paradigm contrast. Both parcellations were created in a data-driven way by decomposing the fMRI data into 22 independent components. We first created a “global” parcellation using functional data from the whole brain as input (top panel). We then selected one parcel from this parcellation that most closely resembles the bilateral auditory cortices (middle panel). We binarised this parcel to create a mask of the auditory cortex. We then decomposed the functional data again using this mask to limit the spatial input to the auditory cortex. This results in a “local” parcellation of the auditory cortex (bottom panel). Please note that we here show the binarised, non-overlapping versions of the parcellations for visualisation purposes. These versions are used to parcellate the paradigm contrast. The original versions of each parcellation are weighted and overlapping and we used these versions to extract time courses from the resting state functional data. **b)** We used ridge regression to predict paradigm activation from rsFC. First, all n participants’ individual rsFC was computed in the two parcellations. The upper triangles of these n FC matrices (each of size Nparcels x Nparcels) were unwrapped and then used as predictor of the vector (of size 1 x Nparcels) containing each parcel’s value in the paradigm contrast.

Since the global parcellation is relatively coarse and might not provide the necessary level of detail to capture individual differences in an auditory oddball paradigm, we created a second, more fine-grained parcellation of the bilateral auditory cortices. It is well established that the auditory cortices are the most relevant regions for an auditory oddball paradigm such as the one we used here (Opitz et al., 1999; Opitz et al., 2002; Schönwiesner et al., 2007; Wible et al., 2001). We refer to this parcellation as “local” parcellation. To be able to compare the two parcellations, we used the same data-driven approach as for the global parcellation. Rather than using an anatomically chosen mask of the auditory cortices, we thus chose the component from the global parcellation that had the closest correspondence to the auditory cortices (shown in **Figure** 1a, centre). We then ran group-level ICA on all participants’ concatenated data only within this auditory cortex mask. This process is summarised in **Figure** 1a. Similar to the global parcellation, we also decomposed the auditory cortex functional data into 25 components, three of which were removed as most closely resembling artefacts. The final auditory cortex sub-parcellation is shown in **Figure** 1a (bottom panel). As for the global parcellation, we plot only the binarised, non-overlapping version of the parcellation as used to parcellate the paradigm contrast. Our approach of creating a local data-driven functional parcellation from a global data-driven functional parcel using ICA is similar to the approach taken in van Oort et al. (2018). The main difference is that in our approach, the time course of the global parcel is not explicitly modelled in the local parcellation, so that local parcels may still contain the time course of the global parcel they are contained within.

### Resting state functional connectivity and paradigm contrast

To compute rsFC, we first extracted time courses within each parcellation from the resting state scanning session. We used dual regression (Beckmann et al., 2009) to extract participant-specific time-courses of the global and local group-ICA parcellations. Dual regression consists of a first step, where the group-level weighted parcels are regressed against each single participant’s voxel time courses as spatial regressors using multiple regression to obtain participant-specific IC time courses; and a second step, in which these participant-specific time courses are used as temporal regressors to obtain participant-specific weights of spatial maps of each parcel. The participant-specific time courses are used to compute rsFC. We used FSLnets (Smith, Vidaurre, et al., 2013) to first standardise the time series and then use Pearson’s correlation to compute the pairwise FC between all parcels. The correlations were then z-transformed. This obtained a matrix of size *N_parcels_* x *N_parcels_* for each participant, in which each element represents the pairwise rsFC between two parcels. Here, *N_participants_* (the number of participants) is 52 and *N_parcels_* (the number of parcels in each parcellation) is 22. We applied the same procedure to both the global and the local parcellation.

The paradigm data was analysed as described in Lumaca, Dietz, et al. (2021) using SPM12. First-level analysis consisted of a generalised linear model (GLM) with standard (STD; implicitly modelled), deviant contour (DC), and deviant interval (DI) regressors convolved with a canonical hemodynamic response function (HRF), and realignment parameters to account for head motion. We contrasted the standards with both types of deviants. We then parcellated the contrast map of each individual into the same parcellations as used for the resting state data. Since the paradigm contrast of a participant consists of only one time point, this could not be done using dual regression as for the resting state data. Instead, the parcellations were first binarised and overlap between parcels was removed. We then extracted values within each parcel as the mean of all voxels belonging to that specific parcel. This resulted in a vector of size 1 x *N_parcels_* for each participant, where the vector elements are summaries of the paradigm contrast value within a given parcel. We extracted values of the paradigm contrast for both the global and the local parcellation.

### Predictive model

To predict individual paradigm activation across the brain, we used a separate ridge regression model for each parcel of the global and the local parcellation, respectively. The general idea of the predictive model is illustrated in **Figure** 1b.

For each model, we used the rsFC of all participants as a predictor. We first reshaped the 3D tensor containing all participants’ rsFC matrices of size *N_participants_* x *N_parcels_* x *N_parcels_* by vectorising the upper triangle of each participant’s *N_parcels_* x *N_parcels_* FC matrix, resulting in a matrix of *N_participants_* x (*N_parcels_* * (*N_parcels_*-1) / 2). We then standardised this matrix column-wise. The outcome variable of each model is the *N_participants_* x 1 vector containing individual contrast values within one parcel. We thus fitted *N_parcels_* (independent) models for each parcellation. In the runs where the outcome variable was paradigm activation in the global parcellation, rsFC in the global parcellation was the predictor, and in the runs where we predicted paradigm activation in the local parcellation, rsFC in the local parcellation was the predictor.

The regression problem is defined as:

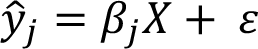

where *ŷ_j_* is the *N_participants_* x 1 vector of predicted values in region *j*, *X* are the features from the rsFC matrix, *β_j_* is the regression parameter for parcel *j* and *ε* is the error (residual) term. The parameter *β_j_* is estimated by minimising:

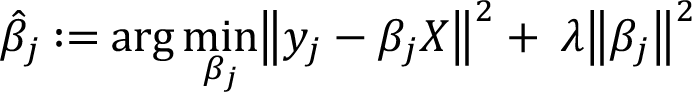

Where *β̂_j_* is the new estimate for *β_j_* and *λ* is a regularisation parameter.

We used 4-fold nested cross-validation to optimise the regularisation parameter (inner loop) and evaluate the accuracy on the test set. We performed 100 iterations of this process where participants were randomly assigned to the different cross-validation folds. This means that the model found slightly different parameters in each iteration and the accuracy therefore varies.

We evaluated the model’s accuracy in each of these iterations in two ways: First, to measure the accuracy of full map predictions, we correlated the model-predicted values *ŷ^i^*with the actual values *y^i^* of a given participant *i* using Pearson’s correlation. This indicates how similar a participant’s predicted values over all parcels are to their actual values. A model with high participant-specificity should produce predicted values that are more similar to the participant’s own actual values than to all other participants’ actual values. We thus compared *r_self_*, i.e., the correlation coefficient between *ŷ^i^* and *y^i^*, with *r_other_*, i.e., the correlation coefficient between *ŷ^i^* and *y^I\{i}^* (*I\{i}* indicating the set of participants not including *i*). Second, to evaluate prediction accuracy within a given parcel *j*, we correlated the model-predicted values *ŷ_j_* with the actual values *y_j_*. This indicates how well the model’s prediction of differences between participants in a specific parcel corresponds to the actual variability in this parcel. We used this measure as a basis to identify parcels that can be better predicted than others, as described in detail in the next section.

The predictive model and following analyses were carried out in Matlab (MATLAB, 2016). Code for ridge regression from FC is available under https://github.com/vidaurre/NetsPredict.

### Parcel-specific probabilistic index

We calculated a parcel-specific probabilistic index to identify parcels where prediction accuracy was unexpectedly high as compared to all other runs and parcels. Our approach is inspired by the probabilistic indices of abnormality introduced by Marquand et al. (2016). Compared to the approach taken in Marquand et al. (2016), we here aimed to find parcels which are positive outliers in terms of model accuracy with respect to the other parcels and runs, rather than identifying participants and brain regions that can be considered abnormal with respect to a normative population. The probabilistic index is therefore calculated for the parcel-level correlation coefficients (correlation between *ŷ_j_* and *y_j_*) instead of the activation values themselves.

The cross-validation folds used to estimate the model’s parameters are randomised 100 times, meaning that we have 100 slightly different iterations of each parcel model. We then calculate the correlation between *ŷ_j_* and *y_j_* for each parcel *j* and each iteration. Next, we fit a normal distribution and the corresponding probability density function (PDF) to all correlation coefficients. Given this function, we can obtain the probability of the correlation coefficient being this particular value or higher by computing the integral of the curve between each obtained correlation coefficient and ∞. We first focus on the right tail of this distribution, containing all values where this probability is smaller than 5% (corresponding to a z-score greater than 1.96). In other words, we are looking at the top 5% of all predictions of between-participant variability in all parcels and runs. We then plot in the brain how many of the 100 iterations of each parcel appeared in this right tail of the distribution using Connectome Workbench (Marcus et al., 2011).

Although this measure may be an interesting indicator of prediction accuracy differences between parcels, the random iterations of the model may only coincidentally appear in the tail of this distribution. We therefore next computed a probabilistic index based on the robust best runs of each parcel. We first computed the z-scores of the obtained correlation coefficient for each parcel and each run with respect to all runs and parcels. We then considered the “block maxima” by calculating the 90% trimmed mean of the highest 10% z-scores within each parcel. In other words, we here focused on the (trimmed) right tail of the distribution of each parcel. These block maxima can be described using an extreme value distribution (EVD) and its corresponding PDF. As before, we can calculate the probability of the block max. z-score of a given parcel being the obtained value or higher within the EVD. The resulting probability value given the EVD of block maxima is a robust, network-specific probabilistic index. This index can be plotted in the brain to identify parcels that are positive outliers in prediction accuracy.

## Results

### Within the auditory cortex, individual task activation can be best predicted in task-relevant right superior temporal gyrus

In the local auditory cortex parcellation, the parcel corresponding to the right superior temporal gyrus (STG) was the best predicted. The next best predicted regions included the left superior temporal gyrus and the posterior part of the right insula. The maps of the best predicted regions within the auditory cortex are shown in Figure 2a.

**Figure 2.**
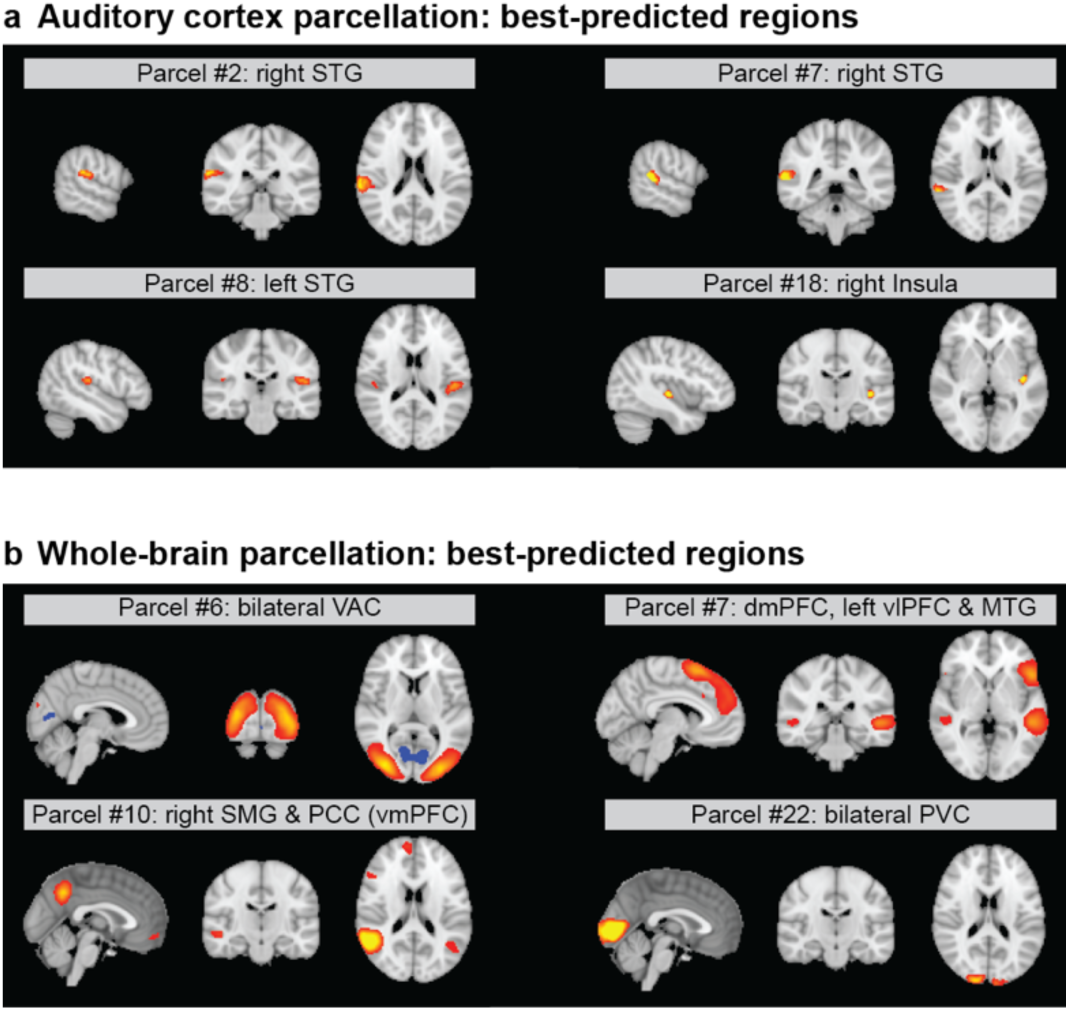
Maps of best predicted regions. The maps show the voxel weights for the components that were best predicted within each parcellation. a) Best predicted regions within auditory cortex parcellation. b) Best predicted regions within whole-brain parcellation. Abbreviations: STG: superior temporal gyrus; VAC: visual association cortex; dmPFC: dorsomedial prefrontal cortex; vlPFC: ventrolateral prefrontal cortex; MTG: middle temporal gyrus; SMG: supramarginal gyrus; PCC: posterior cingulate cortex; vmPFC: ventromedial prefrontal cortex; PVC: primary visual cortex.

We ran each model 100 times, randomly re-assigning subjects to cross-validation folds at each iteration. We then considered where each parcel lies in the distribution of prediction accuracies across all iterations and parcels. Figure 3a shows the distribution of all prediction accuracies, as measured by the correlation coefficient between the predicted interindividual variability and the actual interindividual variability. To estimate the probability of obtaining a certain prediction accuracy, we fitted a Gaussian distribution to the observed prediction accuracy values and considered the probability given the probability density function (PDF) of this distribution. Since we were interested in regions where differences between individuals are significantly better predicted than others, we focused on the right tail of this distribution, where the probability of correlation coefficients *r* being equal to or higher than *r(j,n)* for parcel *j* and run *n* is less than 5%. The dashed line represents the 95^th^ percentile, i.e., where the probability of obtaining a prediction accuracy greater than or equal to the one measured is 5%, corresponding to a z-score of 1.96. We then looked at these results by parcel and run to understand whether all parcels appear equally often in these top 5% of predictions, or whether a small number of parcels appear consistently over runs. Figure 3b plots these probabilities by parcel and run, indicating that the best predictions are not randomly distributed across parcels but that certain parcels consistently appear in the top 5% of predictions, while others never appear. Projecting these counts for each parcel into the brain (see Fig. 3c), we can observe that the region with the highest number of appearances in the top 5% of predictions is the right STG (here parcel #2), which scores in the 95^th^ percentile in 87 out of 100 runs. Only parcels that appear at least once in this tail of the distribution are highlighted.

**Figure 3.**
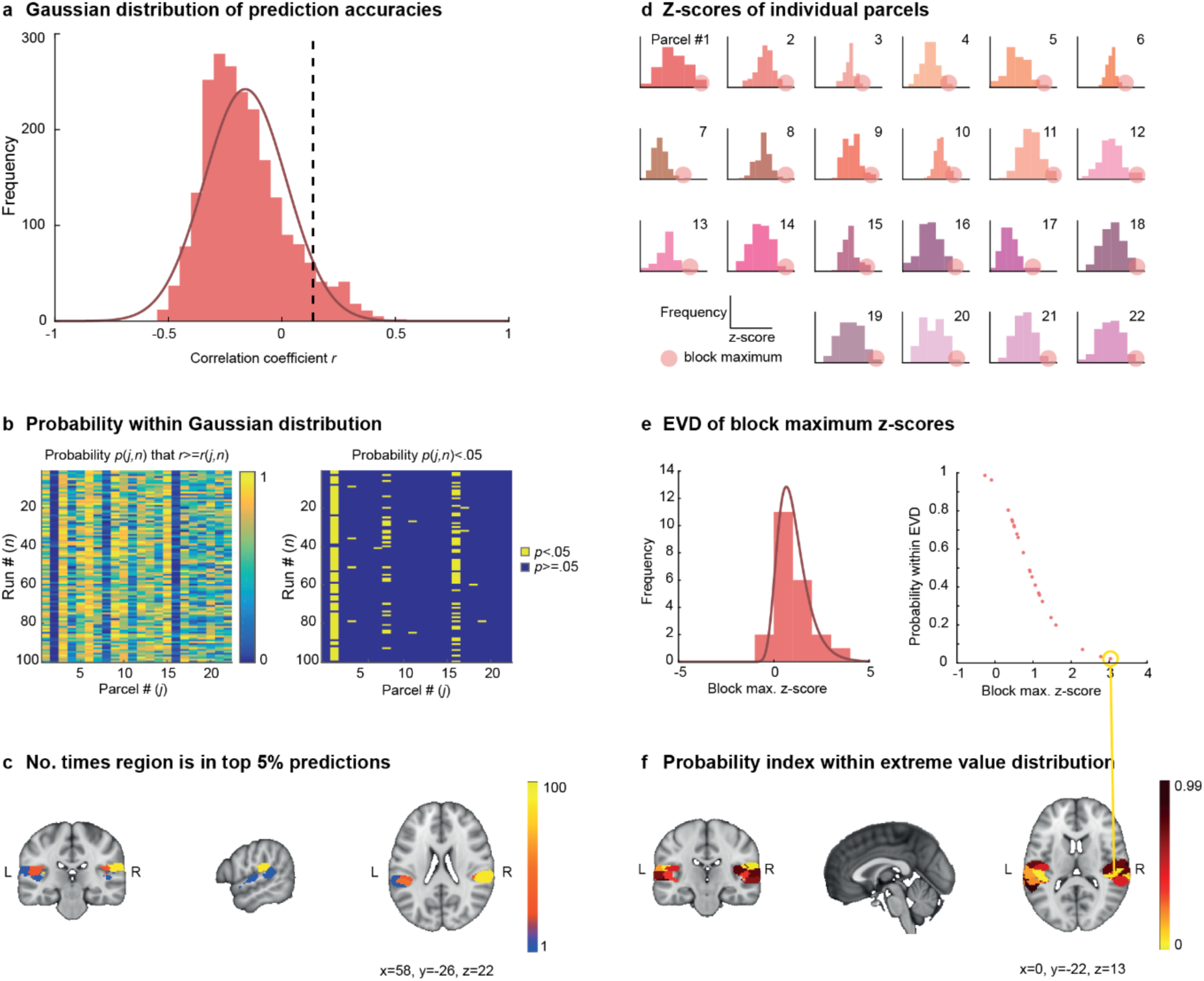
Task activation is robustly better predicted in task-relevant regions of the auditory cortex. **a)** Histogram of prediction accuracies across all 22 parcels and 100 iterations and fitted Gaussian distribution. The dashed line represents the 95^th^ percentile, i.e., the top 5% of predictions. **b)** Probability of obtained prediction accuracy by parcel and run. **c)** Regions that appear more than once in the top 5% predictions within the auditory cortex. **d)** Z-scores and block maxima (the 10% best predicted runs) across runs for each parcel. **e)** Histogram and fitted extreme value distribution (EVD) for 90% trimmed mean block maximum z-scores from d). **f)** Probability index, i.e., probability of obtaining the block max. z-score given the PDF of the EVD from e) across regions in the auditory cortex parcellation. Abbreviations: EVD: extreme value distribution.

Since the cross-validation step introduces randomness in the model and its outcome, single runs of the model are not necessarily a reliable indicator of model performance. To account for this randomness, we next computed a robust, probabilistic index given the best runs within each parcel. Specifically, we computed the 90% trimmed mean of the highest 10% z-scores within each parcel (“block maximum”, see Fig. 3d). We then fitted an extreme value distribution (EVD) to these block maxima (Fig. 3e). Given the PDF of this EVD, we can compute a probabilistic index for each IC, indicating the probability within the EVD that the block max. z-score of a given region would be the obtained value or higher. This allows identifying regions that are positive outliers in terms of prediction accuracy. The relationship between block max. z-scores and the probability within the EVD is shown in 3d-e. In the auditory cortex parcellation, the strongest outlier (i.e., the parcel that is robustly best predicted) is again the right STG (parcel #2) with a block max. z-score of 3.03 (*p*=0.02). The probabilistic index of each region in the brain is plotted in Fig. 3f. Low values indicate that the highest obtained prediction accuracy is significantly higher than the highest obtained prediction accuracy in other parcels.

### Across the brain, visual rather than auditory cortex activation can be best predicted

In the global parcellation, we found the bilateral visual association cortices to be the best predicted. The next best predicted parcels contain the bilateral primary visual cortices, the right primary motor cortex, the posterior cingulate cortex, the bilateral hippocampus, the bilateral caudate nuclei, the medial prefrontal cortex, and the pars triangularis of the right inferior frontal gyrus.

As for the local auditory cortex parcellation, we predicted interindividual differences in task activation for each parcel in the whole-brain parcellation, and we ran each model 100 times. Figure 4a shows the histogram and fitted Gaussian distribution of the obtained prediction accuracies (correlation coefficients between predicted and true interindividual variability) across all parcels and runs, as well as the dashed line cut-off for the 95^th^ percentile. Plotting the probability of the obtained prediction accuracies by parcel and run (Fig. 3b) shows that certain regions appear several times in the top 5% predictions, while other regions never appear. In the whole-brain map, the region with the highest number of appearances in the 95^th^ percentile is the bilateral visual association cortex (here parcel #6) with 55 out of 100 runs.

**Figure 4.**
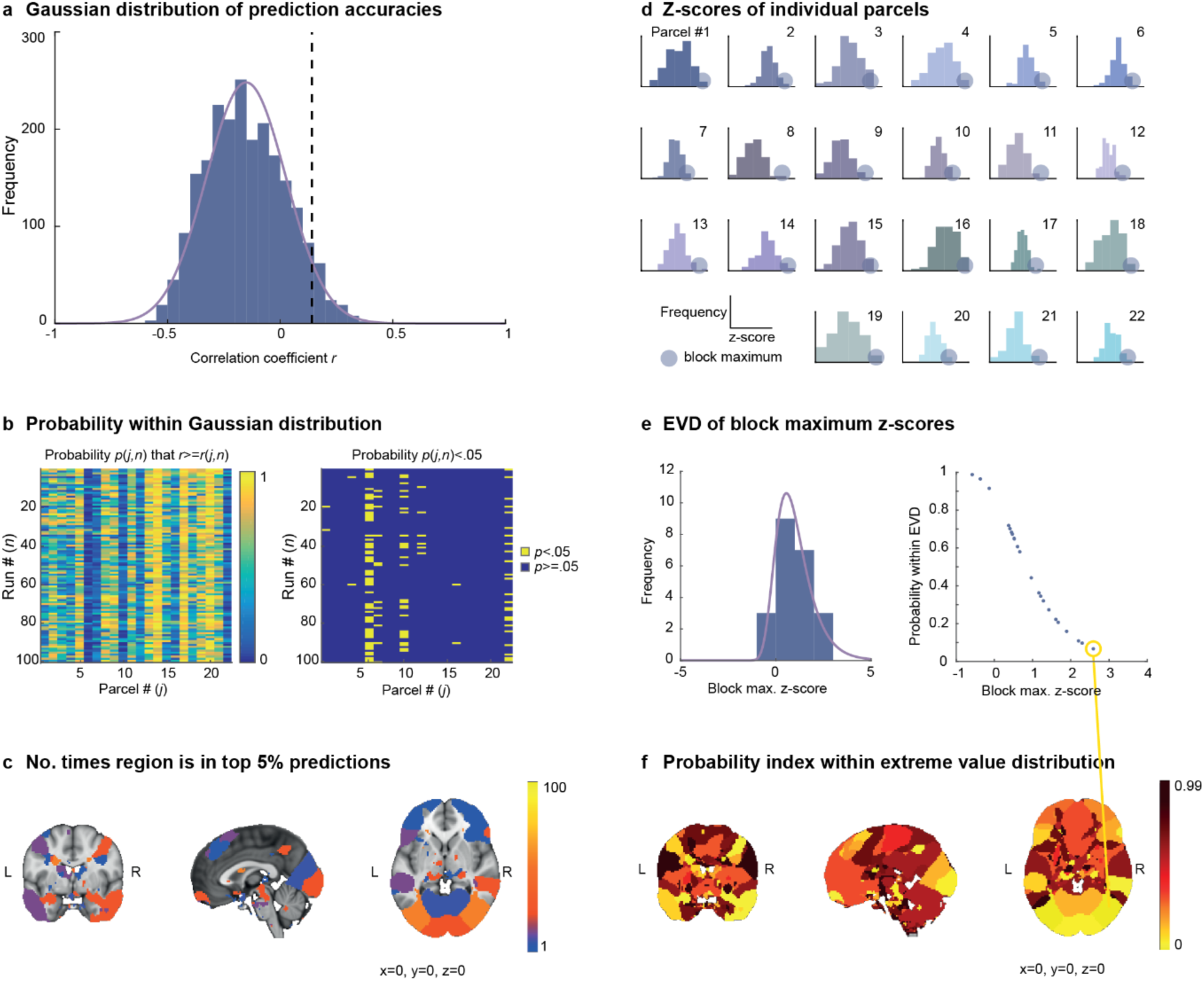
Across the whole brain, task activation is robustly better predicted in visual areas. **a)** Histogram of prediction accuracies across all 22 parcels and 100 iterations and fitted Gaussian distribution. The dashed line represents the 95^th^ percentile, i.e., the top 5% of predictions. **b)** Probability of obtained prediction accuracy by parcel and run. **c)** Regions that appear more than once in the top 5% predictions across the whole brain. **d)** Z-scores and block maxima (the 10% best predicted runs) across runs for each parcel. **e)** Histogram and fitted extreme value distribution (EVD) for 90% trimmed mean block maximum z-scores from d). **f)** Probability index, i.e., probability of obtaining the block max. z-score given the PDF of the EVD from e) across regions in the whole-brain parcellation. Abbreviations: EVD: extreme value distribution.

We then considered, as for the local auditory cortex parcellation, the robust probabilistic index of the best predictions in each parcel. In the whole-brain parcellation, the largest outlier (i.e., the parcel that is robustly best predicted) was again the bilateral visual association cortex (parcel #6) with a block max. z-score of 2.59 (*p*=0.06).

## Discussion

In this study, we aimed at predicting differences between individuals in brain activation patterns during melodic contour processing from their rsFC. We found that individual patterns of paradigm activation were significantly better predicted in some regions than in others. The best predicted parcels were sensory association cortices (visual and auditory), followed by primary sensory cortices (visual and motor), and areas belonging to the default mode network (medial prefrontal cortex, posterior cingulate, hippocampus). In these regions, interindividual differences in paradigm activation between individuals could be accurately predicted, and the superiority of prediction accuracy in these regions was robust across randomised runs of the model.

One reason we expected individual differences to be more predictable in some parcels than in others is that not all regions are relevant for the paradigm at hand. The paradigm we employed specifically requires the processing of melodic contour, a higher-level feature of auditory stimuli concerning the direction of a sequence of tones while disregarding the size of their pitch interval. In the local (auditory cortex) parcellation, we found that individual differences in paradigm activation could be best predicted in the right STG, a part of the auditory association cortices. This area is crucial for the processing of complex auditory representations, such as melodic contour: While more basic auditory features, such as pitch, are processed mainly in the primary auditory cortex, more abstract and complex melodic features are processed in areas further away from the primary auditory cortex, like the STG and parieto-temporal junction (Stewart et al., 2006) (for a review, see Deouell et al. (2007)). Thus, it appears that complex melodic processing is organized hierarchically, with higher-level auditory regions involved in the recognition of melodic directionality (Patterson et al., 2002; Warren & Griffiths, 2003). Brain areas involved in the processing of melodic contour have recently been investigated using multivariate pattern analysis (MVPA). These studies have found that regions in the right (Lee et al., 2011) or bilateral STG and superior temporal sulcus (Schindler et al., 2013) play a role in the processing of ascending or descending melodic contour by showing that local patterns of activity can accurately discriminate between contour categories. These findings are supported by lesion studies showing that lesions in the right superior temporal lobe impair the ability to discriminate the direction of pitch intervals, which forms the basis for melodic contour processing (Johnsrude et al., 2000). Additionally, only right-lateralised lesions appear to cause deficits in melodic contour processing, while lesions in the left hemisphere do not affect this capacity (Peretz, 1990).

Behavioural work also supports the existence of a right-hemisphere bias for the processing of melodic contour. In one experiment, McKinnon and Schellenberg (1997) presented listeners with series of five tones in one ear and instructed them to make a forced-choice judgment regarding the sequence’s contour. Results indicated that performance was better when the sequence was presented to the left ear (i.e., first processed in the right hemisphere) as opposed to the right ear (left hemisphere). This provides evidence for the notion that the right hemisphere has an inclination to process melodic contour. The involvement of right STG/STS in the processing of contour features is also supported by neuroimaging work on language processing (Kyong et al., 2014). Right superior temporal regions exhibited the largest activity for spoken language with modulation of suprasegmental contour features. Our finding that the right STG can be better predicted than other parts of the auditory cortices may thus be due to this area’s relevance to processing melodic contour, a central requirement of the paradigm.

The second reason for differences between regions in predictability is that activation patterns in some brain regions may be more individual-specific than in others. Several studies aiming to find individual differences in paradigm activation have found sensory (association) cortices to be best predicted, despite not being expected to be relevant to the paradigm at hand.

Villalta-Gil et al. (2017) investigated individual responses to fear paradigms, expecting to find correlations with activation in the amygdala, but found them reflected in the visual cortex instead. When averaging over 47 different paradigm contrasts, Lacosse et al. (2021) found individual differences in paradigm activation to be most predictable in the sensory (visual, auditory, and motor) association cortices. Similarly, Mueller et al. (2013) found that individual variability in FC is higher in heteromodal association cortices compared to unimodal sensory cortices and that brain regions that have higher between-participant variability in FC are better predictors of behaviour in several cognitive domains. Here, using an auditory paradigm with concurrent visual stimuli, we also found the visual association cortices to be best predicted in the global parcellation. Within the local auditory cortex parcellation, the best predicted area is the right STG, which is considered to be an auditory association area (Zevin, 2009). A recent study also investigated inter-individual variability based on intrinsic rsFC and a passive listening paradigm in different sites within the auditory cortices in humans and macaques (Ren et al., 2021) and found the non-primary auditory cortices to be the most distinct. In the brain, phylogenetically old regions like the brain stem and deep subcortical regions that serve fundamental survival functions may be more similar between individuals, while higher-order areas in the neocortex may exhibit differences according to an individual’s unique traits. Indeed, fibre connections in frontal and limbic areas differ little between individuals, while temporal and occipital regions are very diverse (Kerepesi et al., 2018). Inter-individual variability of functional connectivity is larger in frontal and parietal cortices, phylogenetically and ontogenetically late-developing regions (Kaas, 2006; Smaers et al., 2011; Van Essen & Dierker, 2007), whose protracted plasticity would expose them longer to experiential factors (Petanjek et al., 2011). Both paradigm-relevance and a more general level of information about the individual that is intrinsic to sensory association areas may therefore influence predictability across brain regions.

While the prediction of individual differences in paradigm activation from rsFC has the potential to make an important link between fingerprinting studies and the understanding of cognitive function, it seems that more work is needed before this method can be reliably used across the brain in real-life fMRI datasets. An evident limitation is the number of participants necessary for the estimation of these models. Although the sample in this study (*N*=52) is large for an fMRI study, the optimal sample size may be more likely in the order of hundreds, such as in most of the studies previously reported (Cohen et al., 2020; Cole et al., 2016; Parker Jones et al., 2016; Tavor et al., 2016). However, even using a sample size of *N*=200, a recent replication study found that most methods that have been reported to be able to predict paradigm activation maps from rsFC are optimistic and in fact do not outperform simple baseline models (Lacosse et al., 2021). Improving these modelling efforts is an active area of research. It is possible, however, that the difficulty of predicting paradigm activation from rsFC lies in the nature of the data rather than in the details of the models.

While both structural connectivity and rsFC naturally vary greatly between participants, classic fMRI paradigms are designed to constrain brain activity to the studied function. Since the goal of these paradigms is to elucidate brain activity underlying a specific cognitive function by averaging across the group, they allow little variance between participants by definition. Large enough variance between participants is a fundamental requirement for a predictive model to perform well. Lacosse et al. (2021) also found in their study that only one of the 47 contrasts studied could reliably be predicted. It is possible that more naturalistic, unconstrained paradigms could be more easily predicted than the ones studied here and in previous studies.

Modelling individual paradigm activation patterns from rsFC is a worthy goal, since it may be of great clinical relevance for patients unable to complete a certain task. It has, for instance been shown that it is possible to use predicted individual activity patterns to personalise surgical targets (Niu et al., 2021; Parker Jones et al., 2016).

In sum, we here show that differences between individuals in an auditory oddball paradigm of melodic contour can be best predicted in sensory association areas. Creating a sub-parcellation of the auditory cortices made it possible to identify the right STG, an area that is both relevant to the paradigm and likely carries a high amount of individual information, as a site where differences between individuals in paradigm activation can be predicted well. The prediction of individual differences in a paradigm from rsFC depends on each brain area’s relevance to the paradigm and their level of inter-individual variability. While activation patterns in some areas may thus be similar across individuals, sensory association cortices are unique - akin to a fingerprint.

## References

1. Bailes, F. (2010). Dynamic melody recognition: Distinctiveness and the role of musical expertise. Memory & Cognition, 38(5), 641–650. 10.3758/MC.38.5.641

2. Beckmann, C. F. (2012). Modelling with independent components. Neuroimage, 62(2), 891–901. 10.1016/j.neuroimage.2012.02.020

3. Beckmann, C. F., Mackay, C. E., Filippini, N., & Smith, S. M. (2009). Group comparison of resting-state FMRI data using multi-participant ICA and dual regression. Neuroimage, 47, S148. 10.1016/S1053-8119(09)71511-3

4. Biswal, B. B., Mennes, M., Zuo, X. N., Gohel, S., Kelly, C., Smith, S. M., Beckmann, C. F., Adelstein, J. S., Buckner, R. L., Colcombe, S., Dogonowski, A. M., Ernst, M., Fair, D., Hampson, M., Hoptman, M. J., Hyde, J. S., Kiviniemi, V. J., Kötter, R., Li, S. J., . . . Milham, M. P. (2010). Toward discovery science of human brain function [Article]. Proceedings of the National Academy of Sciences of the United States of America, 107(10), 4734–4739. 10.1073/pnas.0911855107

5. Cohen, A. D., Chen, Z., Parker Jones, O., Niu, C., & Wang, Y. (2020). Regression-based machine-learning approaches to predict task activation using resting-state fMRI. Human Brain Mapping, 41(3), 815–826. 10.1002/hbm.24841

6. Cole, M. W., Ito, T., Bassett, D. S., & Schultz, D. H. (2016). Activity flow over resting-state networks shapes cognitive task activations. Nat Neurosci, 19(12), 1718–1726. 10.1038/nn.4406

7. Deouell, L. Y., Deutsch, D., Scabini, D., Soroker, N., & Knight, R. T. (2007). No disillusions in auditory extinction: perceiving a melody comprised of unperceived notes. Front Hum Neurosci, 1, 15. 10.3389/neuro.09.015.2007

8. Dowling, W. J., & Fujitani, D. S. (1971). Contour, interval, and pitch recognition in memory for melodies. Journal of the Acoustical Society of America, 49, 524–531. 10.1121/1.1912382

9. Finn, E. S., Shen, X., Scheinost, D., Rosenberg, M. D., Huang, J., Chun, M. M., Papademetris, X., & Constable, R. T. (2015). Functional connectome fingerprinting: Identifying individuals using patterns of brain connectivity. Nature Neuroscience, 18(11), 1664–1671. 10.1038/nn.4135

10. Fujioka, T., Trainor, L. J., Ross, B., Kakigi, R., & Pantev, C. (2004). Musical Training Enhances Automatic Encoding of Melodic Contour and Interval Structure. Journal of Cognitive Neuroscience, 16(6), 1010–1021. 10.1162/0898929041502706

11. Granier-Deferre, C., Bassereau, S., Ribeiro, A., Jacquet, A.-Y., & DeCasper, A. J. (2011). A Melodic Contour Repeatedly Experienced by Human Near-Term Fetuses Elicits a Profound Cardiac Reaction One Month after Birth. PLOS ONE, 6(2), e17304. 10.1371/journal.pone.0017304

12. Griffanti, L., Douaud, G., Bijsterbosch, J., Evangelisti, S., Alfaro-Almagro, F., Glasser, M. F., Duff, E. P., Fitzgibbon, S., Westphal, R., Carone, D., Beckmann, C. F., & Smith, S. M. (2017). Hand classification of fMRI ICA noise components. Neuroimage, 154, 188–205. 10.1016/j.neuroimage.2016.12.036

13. Griffanti, L., Salimi-Khorshidi, G., Beckmann, C. F., Auerbach, E. J., Douaud, G., Sexton, C. E., Zsoldos, E., Ebmeier, K. P., Filippini, N., Mackay, C. E., Moeller, S., Xu, J., Yacoub, E., Baselli, G., Ugurbil, K., Miller, K. L., & Smith, S. M. (2014). ICA-based artefact removal and accelerated fMRI acquisition for improved resting state network imaging. Neuroimage, 95, 232–247. 10.1016/j.neuroimage.2014.03.034

14. Halpern, A. R., Bartlett, J. C., & Dowling, W. J. (1998). Perception of Mode, Rhythm, and Contour in Unfamiliar Melodies: Effects of Age and Experience. Music Perception, 15(4), 335–355. 10.2307/40300862

15. Horien, C., Shen, X., Scheinost, D., & Constable, R. T. (2019). The individual functional connectome is unique and stable over months to years. Neuroimage, 189, 676–687. 10.1016/j.neuroimage.2019.02.002

16. Jenkinson, M., Beckmann, C. F., Behrens, T. E., Woolrich, M. W., & Smith, S. M. (2012). FSL. Neuroimage, 62(2), 782–790. 10.1016/j.neuroimage.2011.09.015

17. Jeong, E., & Ryu, H. (2016). Melodic Contour Identification Reflects the Cognitive Threshold of Aging. Frontiers in aging neuroscience, 8, 134–134. 10.3389/fnagi.2016.00134

18. Johnsrude, I. S., Penhune, V. B., & Zatorre, R. J. (2000). Functional specificity in the right human auditory cortex for perceiving pitch direction. Brain, 123(1), 155–163. 10.1093/brain/123.1.155

19. Kerepesi, C., Szalkai, B., Varga, B., & Grolmusz, V. (2018). Comparative connectomics: Mapping the inter-individual variability of connections within the regions of the human brain. Neuroscience Letters, 662, 17–21. 10.1016/j.neulet.2017.10.003

20. Kraus, B. T., Perez, D., Ladwig, Z., Seitzman, B. A., Dworetsky, A., Petersen, S. E., & Gratton, C. (2021). Network variants are similar between task and rest states. NeuroImage, 229, Article 117743. 10.1016/j.neuroimage.2021.117743

21. Krienen, F. M., Yeo, B. T., & Buckner, R. L. (2014). Reconfigurable task-dependent functional coupling modes cluster around a core functional architecture. Philos Trans R Soc Lond B Biol Sci, 369(1653). 10.1098/rstb.2013.0526

22. Kyong, J. S., Scott, S. K., Rosen, S., Howe, T. B., Agnew, Z. K., & McGettigan, C. (2014). Exploring the roles of spectral detail and intonation contour in speech intelligibility: an FMRI study. J Cogn Neurosci, 26(8), 1748–1763. 10.1162/jocn_a_00583

23. Kaas, J. H. (2006). Evolution of the neocortex. Curr Biol, 16(21), R910–914. 10.1016/j.cub.2006.09.057

24. Lacosse, E., Scheffler, K., Lohmann, G., & Martius, G. (2021). Jumping over baselines with new methods to predict activation maps from resting-state fMRI. Scientific Reports, 11(1), Article 3480. 10.1038/s41598-021-82681-8

25. Langs, G., Golland, P., & Ghosh, S. S. (2015). Predicting Activation Across Individuals with Resting-State Functional Connectivity Based Multi-Atlas Label Fusion. Medical Image Computing and Computer-Assisted Intervention -- MICCAI 2015, Cham.

26. Lee, Y.-S., Janata, P., Frost, C., Hanke, M., & Granger, R. (2011). Investigation of melodic contour processing in the brain using multivariate pattern-based fMRI. NeuroImage, 57(1), 293–300. 10.1016/j.neuroimage.2011.02.006

27. Lo, C. Y., McMahon, C. M., Looi, V., & Thompson, W. F. (2015). Melodic Contour Training and Its Effect on Speech in Noise, Consonant Discrimination, and Prosody Perception for Cochlear Implant Recipients. Behavioural Neurology, 2015, 352869. 10.1155/2015/352869

28. Lumaca, M., Baggio, G., & Vuust, P. (2021). White matter variability in auditory callosal pathways contributes to variation in the cultural transmission of auditory symbolic systems. Brain Struct Funct, 226(6), 1943–1959. 10.1007/s00429-021-02302-y

29. Lumaca, M., Dietz, M. J., Hansen, N. C., Quiroga-Martinez, D. R., & Vuust, P. (2021). Perceptual learning of tone patterns changes the effective connectivity between Heschl’s gyrus and planum temporale. Human Brain Mapping, 42(4), 941–952. 10.1002/hbm.25269

30. Lumaca, M., Kleber, B., Brattico, E., Vuust, P., & Baggio, G. (2019). Functional connectivity in human auditory networks and the origins of variation in the transmission of musical systems. eLife, 8, e48710. 10.7554/eLife.48710

31. Marcus, D., Harwell, J., Olsen, T., Hodge, M., Glasser, M., Prior, F., Jenkinson, M., Laumann, T., Curtiss, S., & Van Essen, D. (2011). Informatics and Data Mining Tools and Strategies for the Human Connectome Project [Technology Report]. Frontiers in Neuroinformatics, 5(4). 10.3389/fninf.2011.00004

32. Marek, S., Tervo-Clemmens, B., Nielsen, A. N., Wheelock, M. D., Miller, R. L., Laumann, T. O., Earl, E., Foran, W. W., Cordova, M., Doyle, O., Perrone, A., Miranda-Dominguez, O., Feczko, E., Sturgeon, D., Graham, A., Hermosillo, R., Snider, K., Galassi, A., Nagel, B. J., . . . Dosenbach, N. U. F. (2019). Identifying reproducible individual differences in childhood functional brain networks: An ABCD study. Developmental Cognitive Neuroscience, 40, 100706. 10.1016/j.dcn.2019.100706

33. Marquand, A. F., Rezek, I., Buitelaar, J., & Beckmann, C. F. (2016). Understanding Heterogeneity in Clinical Cohorts Using Normative Models: Beyond Case-Control Studies. Biological psychiatry, 80(7), 552–561. 10.1016/j.biopsych.2015.12.023

34. MATLAB. (2016). R2016b. The MathWorks Inc.

35. McKinnon, M. C., & Schellenberg, E. G. (1997). A left-ear advantage for forced-choice judgements of melodic contour. Can J Exp Psychol, 51(2), 171–175. 10.1037/1196-1961.51.2.171

36. Mueller, S., Wang, D., Fox, M. D., Yeo, B. T., Sepulcre, J., Sabuncu, M. R., Shafee, R., Lu, J., & Liu, H. (2013). Individual variability in functional connectivity architecture of the human brain. Neuron, 77(3), 586–595. 10.1016/j.neuron.2012.12.028

37. Niu, C., Cohen, A. D., Wen, X., Chen, Z., Lin, P., Liu, X., Menze, B. H., Wiestler, B., Wang, Y., & Zhang, M. (2021). Modeling motor task activation from resting-state fMRI using machine learning in individual participants. Brain Imaging Behav, 15(1), 122–132. 10.1007/s11682-019-00239-9

38. Opitz, B., Mecklinger, A., von Cramon, D. Y., & Kruggel, F. (1999). Combining electrophysiological and hemodynamic measures of the auditory oddball. Psychophysiology, 36(1), 142–147. 10.1017/S0048577299980848

39. Opitz, B., Rinne, T., Mecklinger, A., von Cramon, D. Y., & Schröger, E. (2002). Differential contribution of frontal and temporal cortices to auditory change detection: fMRI and ERP results. NeuroImage, 15(1), 167–174. 10.1006/nimg.2001.0970

40. Osher, D. E., Brissenden, J. A., & Somers, D. C. (2019). Predicting an individual’s dorsal attention network activity from functional connectivity fingerprints. Journal of Neurophysiology, 122(1), 232–240. 10.1152/jn.00174.2019

41. Pankanti, S., Prabhakar, S., & Jain, A. K. (2002). On the Individuality of Fingerprints. IEEE Trans. Pattern Anal. Mach. Intell., 24(8), 1010–1025. 10.1109/tpami.2002.1023799

42. Parker Jones, O., Voets, N. L., Adcock, J. E., Stacey, R., & Jbabdi, S. (2016). Resting connectivity predicts task activation in pre-surgical populations. NeuroImage. Clinical, 13, 378–385. 10.1016/j.nicl.2016.12.028

43. Patterson, R. D., Uppenkamp, S., Johnsrude, I. S., & Griffiths, T. D. (2002). The processing of temporal pitch and melody information in auditory cortex. Neuron, 36(4), 767–776. 10.1016/s0896-6273(02)01060-7

44. Peretz, I. (1990). Processing of local and global musical information by unilateral brain-damaged patients. Brain, 113 (Pt 4), 1185–1205. 10.1093/brain/113.4.1185

45. Peretz, I., & Babaï, M. (1992). The role of contour and intervals in the recognition of melody parts: Evidence from cerebral asymmetries in musicians. Neuropsychologia, 30(3), 277–292. 10.1016/0028-3932(92)90005-7

46. Petanjek, Z., Judaš, M., Šimic, G., Rasin, M. R., Uylings, H. B., Rakic, P., & Kostovic, I. (2011). Extraordinary neoteny of synaptic spines in the human prefrontal cortex. Proc Natl Acad Sci U S A, 108(32), 13281–13286. 10.1073/pnas.1105108108

47. Ren, J., Xu, T., Wang, D., Li, M., Lin, Y., Schoeppe, F., Ramirez, J. S. B., Han, Y., Luan, G., Li, L., Liu, H., & Ahveninen, J. (2021). Individual Variability in Functional Organization of the Human and Monkey Auditory Cortex. Cerebral Cortex, 31(5), 2450–2465. 10.1093/cercor/bhaa366

48. Salimi-Khorshidi, G., Douaud, G., Beckmann, C. F., Glasser, M. F., Griffanti, L., & Smith, S. M. (2014). Automatic denoising of functional MRI data: combining independent component analysis and hierarchical fusion of classifiers. Neuroimage, 90, 449–468. 10.1016/j.neuroimage.2013.11.046

49. Schindler, A., Herdener, M., & Bartels, A. (2013). Coding of Melodic Gestalt in Human Auditory Cortex. Cerebral Cortex, 23(12), 2987–2993. 10.1093/cercor/bhs289

50. Schönwiesner, M., Krumbholz, K., Rübsamen, R., Fink, G. R., & von Cramon, D. Y. (2007). Hemispheric asymmetry for auditory processing in the human auditory brain stem, thalamus, and cortex. Cereb Cortex, 17(2), 492–499. 10.1093/cercor/bhj165

51. Smaers, J. B., Steele, J., Case, C. R., Cowper, A., Amunts, K., & Zilles, K. (2011). Primate prefrontal cortex evolution: human brains are the extreme of a lateralized ape trend. Brain Behav Evol, 77(2), 67–78. 10.1159/000323671

52. Smith, S. M., Beckmann, C. F., Andersson, J., Auerbach, E. J., Bijsterbosch, J., Douaud, G., Duff, E., Feinberg, D. A., Griffanti, L., Harms, M. P., Kelly, M., Laumann, T., Miller, K. L., Moeller, S., Petersen, S., Power, J., Salimi-Khorshidi, G., Snyder, A. Z., Vu, A. T., . . . Consortium, W. U.-M. H. (2013). Resting-state fMRI in the Human Connectome Project. Neuroimage, 80, 144–168. 10.1016/j.neuroimage.2013.05.039

53. Smith, S. M., Fox, P. T., Miller, K. L., Glahn, D. C., Fox, P. M., Mackay, C. E., Filippini, N., Watkins, K. E., Toro, R., Laird, A. R., & Beckmann, C. F. (2009). Correspondence of the brain’s functional architecture during activation and rest. Proceedings of the National Academy of Sciences, 106(31), 13040–13045. 10.1073/pnas.0905267106

54. Smith, S. M., Vidaurre, D., Beckmann, C. F., Glasser, M. F., Jenkinson, M., Miller, K. L., Nichols, T. E., Robinson, E. C., Salimi-Khorshidi, G., Woolrich, M. W., Barch, D. M., Uğurbil, K., & Van Essen, D. C. (2013). Functional connectomics from resting-state fMRI. Trends in Cognitive Sciences, 17(12), 666–682. 10.1016/j.tics.2013.09.016

55. Stewart, L., von Kriegstein, K., Warren, J. D., & Griffiths, T. D. (2006). Music and the brain: disorders of musical listening. Brain, 129(10), 2533–2553. 10.1093/brain/awl171

56. Tavor, I., Jones, O. P., Mars, R. B., Smith, S. M., Behrens, T. E., & Jbabdi, S. (2016). Task-free MRI predicts individual differences in brain activity during task performance. Science, 352(6282), 216–220. 10.1126/science.aad8127

57. Tobyne, S. M., Somers, D. C., Brissenden, J. A., Michalka, S. W., Noyce, A. L., & Osher, D. E. (2018). Prediction of individualized task activation in sensory modality-selective frontal cortex with ’connectome fingerprinting’. NeuroImage, 183, 173–185. 10.1016/j.neuroimage.2018.08.007

58. Van Essen, D. C., & Dierker, D. L. (2007). Surface-based and probabilistic atlases of primate cerebral cortex. Neuron, 56(2), 209–225. 10.1016/j.neuron.2007.10.015

59. van Oort, E. S. B., Mennes, M., Navarro Schröder, T., Kumar, V. J., Zaragoza Jimenez, N. I., Grodd, W., Doeller, C. F., & Beckmann, C. F. (2018). Functional parcellation using time courses of instantaneous connectivity. Neuroimage, 170, 31–40. 10.1016/j.neuroimage.2017.07.027

60. Villalta-Gil, V., Hinton, K. E., Landman, B. A., Yvernault, B. C., Perkins, S. F., Katsantonis, A. S., Sellani, C. L., Lahey, B. B., & Zald, D. H. (2017). Convergent individual differences in visual cortices, but not the amygdala across standard amygdalar fMRI probe tasks. NeuroImage, 146, 312–319. 10.1016/j.neuroimage.2016.11.038

61. Warren, J. D., & Griffiths, T. D. (2003). Distinct Mechanisms for Processing Spatial Sequences and Pitch Sequences in the Human Auditory Brain. The Journal of Neuroscience, 23(13), 5799–5804. 10.1523/jneurosci.23-13-05799.2003

62. Wible, C. G., Kubicki, M., Yoo, S. S., Kacher, D. F., Salisbury, D. F., Anderson, M. C., Shenton, M. E., Hirayasu, Y., Kikinis, R., Jolesz, F. A., & McCarley, R. W. (2001). A functional magnetic resonance imaging study of auditory mismatch in schizophrenia. The American journal of psychiatry, 158(6), 938–943. 10.1176/appi.ajp.158.6.938

63. Zevin, J. (2009). Word Recognition. In L. R. Squire (Ed.), Encyclopedia of Neuroscience (pp. 517–522). Oxford Academic Press. 10.1016/B978-008045046-9.01881-7

